# Cross-neutralization activity against SARS-CoV-2 is present in currently available intravenous immunoglobulins

**DOI:** 10.1101/2020.06.19.160879

**Authors:** José-María Díez, Carolina Romero, Júlia Vergara-Alert, Melissa Belló-Perez, Jordi Rodon, José Manuel Honrubia, Joaquim Segalés, Isabel Sola, Luis Enjuanes, Rodrigo Gajardo

## Abstract

**Background:** There is a crucial need for effective therapies that are immediately available to counteract COVID-19 disease. Recently, ELISA binding cross-reactivity against components of human epidemic coronaviruses with currently available intravenous immunoglobulins (IVIG) Gamunex-C and Flebogamma DIF (5% and 10%) have been reported. In this study, the same products were tested for neutralization activity against SARS-CoV-2, SARS-CoV and MERS-CoV and their potential as an antiviral therapy.

**Methods:** The neutralization capacity of six selected lots of IVIG was assessed against SARS-CoV-2 (two different isolates), SARS-CoV and MERS-CoV in cell cultures. Infectivity neutralization was measured by determining the percent reduction in plaque-forming units (PFU) and by cytopathic effects for two IVIG lots in one of the SARS-CoV-2 isolates. Neutralization was quantified using the plaque reduction neutralization test 50 (PRNT_50_) in the PFU assay and the half maximal inhibitory concentration (IC_50_) in the cytopathic/cytotoxic method (calculated as the minus log_10_ dilution which reduced the viral titer by 50%).

**Results:** All IVIG preparations showed neutralization of both SARS-CoV-2 isolates, ranging from 79 to 89.5% with PRNT_50_ titers from 4.5 to >5 for the PFU method and ranging from 47.0%-64.7% with an IC_50_ ~1 for the cytopathic method. All IVIG lots produced neutralization of SARS-CoV ranging from 39.5 to 55.1 % and PRNT_50_ values ranging from 2.0 to 3.3. No IVIG preparation showed significant neutralizing activity against MERS-CoV.

**Conclusion:** In cell culture neutralization assays, the tested IVIG products contain antibodies with significant cross-neutralization capacity against SARS-CoV-2 and SARS-CoV. However, no neutralization capacity was demonstrated against MERS-CoV. These preparations are currently available and may be immediately useful for COVID-19 management.

## Introduction

The outbreak of the novel Severe Acute Respiratory Syndrome Coronavirus 2 (SARS-CoV-2) which causes the respiratory disease COVID-19 was declared a pandemic by the WHO in March 2020. Most infected patients (80%) have mild symptoms. However, about 20% of COVID-19 patients can progress to severe pneumonia and to acute respiratory distress syndrome (ARDS) which is associated with multi-organ failure and death ^1^. The current critical situation demands an effective and reliable therapy that is immediately available to control the progression of the disease. Convalescent plasma or plasma-derived immunoglobulin (IG) (either polyvalent IG prepared from healthy donors or hyperimmune IG prepared from donors with high antibody titers against a specific antigen) have been historically used as a readily available therapeutic option in outbreaks of emerging or re-emerging infections ^2^.

To date, seven human coronaviruses (HCoV) have been identified. Four of them are endemic and globally distributed (HCoV-229E, HCoV-NL63, HCoV-OC43 and HCoV-HKU1) ^3^. These viruses typically cause mild symptoms and are associated with about 15% of common colds ^4^. However, the three other human coronaviruses (SARS-CoV, MERS-CoV and SARS-CoV-2) are zoonotic epidemic viruses that can cause severe respiratory infections and fatalities. Severe acute respiratory syndrome coronavirus (SARS-CoV) emerged in China in 2002 with the last reported case in 2014. Middle East respiratory syndrome coronavirus (MERS-CoV) emerged in Saudi Arabia a decade later, in 2012, and led to an outbreak in South Korea in 2015. MERS-CoV still emerges sporadically in humans from its reservoir in camelids ^5–7^. More recently (December 2019), the novel coronavirus SARS-CoV-2 emerged in China and because of its extraordinary human-to-human transmissibility is currently causing an unprecedented pandemic ^8^. Although several therapeutic approaches against SARS-CoV-2 are under investigation, therapeutic agents of proven efficacy are still lacking. Interestingly, coronaviruses share some morphological and functional properties that may be associated with cross-reactive immune responses. This cross-reactivity may have important therapeutic implications ^9^.

SARS-CoV, SARS-CoV-2, and MERS-CoV are classified within the family *Coronaviridae*, genus *Betacoronavirus*, subgenera *Sarbecovirus* (SARS-CoV, SARS-CoV-2) and *Merbecovirus* (MERS-CoV). The spike protein (S), which is exposed on the virion surface, is the main determinant of the coronavirus entry into the host cell and is also the major target of neutralizing antibodies ^10^. Spikes are formed by trimers of protein S, which is in turn formed by subunit (S1) that mediates the binding to the cell receptor and a membrane-anchored subunit (S2) that mediates the fusion of the virus with cell membranes ^11^. Potent neutralizing antibodies often target the receptor interaction site on S1. However, the S1 subunit shows a higher variability than S2. Antibodies targeting S1 are often virus-specific making S2 a better target for cross-neutralizing antibodies ^12,13^.

The amino-acid sequence identity among the S proteins of human betacoronaviruses causing mild (HCoV-OC43 and HCoV-HKU1) and severe (SARS-CoV, SARS-CoV-2, and MERS-CoV) respiratory infections varies between 22% and 33% ^10^. However, the S proteins of SARS-CoV and SARS-CoV-2 share 77% amino-acid identity ^14^ and more than 90% RNA sequence homology ^15^. Cross-reactivity in antigenic responses has been described among human coronaviruses of the same genus, particularly betacoronaviruses. Cross-reactivity between SARS-CoV, MERS-CoV and other endemic human coronaviruses has been reported in some neutralization assays ^16–18^.

Recently, cross-reactivity in ELISA binding assays against antigens of SARS-CoV, SARS-CoV-2, and MERS-CoV has been reported with currently available intravenous immunoglobulins (IVIG) such as Gamunex-C and Flebogamma DIF ^19^. In this study, the neutralization capacity of the IVIG products Gamunex-C and Flebogamma DIF against these epidemic human coronaviruses ‒SARS-CoV, SARS-CoV-2, and MERS-CoV‒ was evaluated.

## Material and Methods

### Experimental products

IVIG products used in this study were Flebogamma^®^ DIF 5% and 10% (Instituto Grifols S.A., Barcelona, Spain) and Gamunex^®^-C 10% (Grifols Therapeutics Inc., Raleigh NC, USA), two highly purified (≥ 98%-99%IgG), unmodified human immunoglobulins. Each product is manufactured from plasma collected from thousands of donors in the US and/or several European countries. IgG concentrations in Flebogamma DIF products were 50 mg/mL and 100 mg/mL (5% and 10%) and in Gamunex-C, the concentration was 100 mg/mL (10%). To ensure a virus-free product, both IVIG manufacturing processes contain dedicated steps with high pathogen clearance capacity, such as solvent/detergent treatment, heat treatment, caprylate treatment and Planova™ nanofiltration down to 20 nm pore size. The plasma used to manufacture the IVIG lots tested was collected from March 2018 to October 2019.

### Study design

Six different lots of Flebogamma DIF and Gamunex-C were tested at several dilutions for cross-reactivity against SARS-CoV, SARS-CoV-2, and MERS-CoV by: i) ELISA techniques; and ii) well-stablished neutralization assays in cell cultures. Lots were identified as F1 and F2 for Flebogamma 5% DIF, F3 and F4 for Flebogamma 10% DIF and G1 and G2 for Gamunex-C. Each experiment was performed in duplicate. Handling of viruses and cell cultures was carried out at the Level 3 Biosafety Laboratories in the *Centro Nacional de Biotecnología - Consejo Superior de Investigaciones Científicas* (CNB-CSIC; Madrid, Spain) and the *Institut de Recerca i Tecnologia Agroalimentàries - Centre de Recerca en Sanitat Animal* (IRTA-CReSA; Barcelona, Spain), following the centers’ biohazard safety guidelines and under authorizations #A/ES/00/I-8 and #SA-10430-20, respectively.

### Virus strains

Recombinant SARS-CoV was generated from Urbani strain using a previously described reverse genetic technique ^20^. Two different SARS-CoV-2 isolates were tested: a) SARS-CoV-2 MAD6 isolated from a COVID-19 patient in Spain; and b) SARS-CoV-2 (accession ID EPI_ISL_418268 at GISAID repository: http://gisaid.org) isolated from a COVID-19 patient in Spain. Both stock viruses (a and b) were prepared by collecting the supernatant from Vero E6 cells, as previously described ^21^. Recombinant MERS-CoV was generated using a previously described reverse genetic system ^22^ from the reference sequence of MERS-CoV isolated from the index patient EMC/2012 (GeneBank JX869059) ^23^.

### Cell lines and cultures

Huh7 is a well differentiated human hepatocyte-derived carcinoma cell line, kindly provided by Dr. Luis Carrasco (Centr*o de Biología Molecular Severo Ochoa - Consejo Superior de Investigaciones Científicas* [CBMSO-CSIC], Madrid, Spain). Huh7 is composed of epithelial-like cells susceptible to infection by MERS-CoV ^24^.

Vero E6 is a cell line isolated from kidney epithelial cells extracted from an African green monkey. Vero E6 is composed of epithelial-like cells susceptible to infection by SARS-CoV and SARS-CoV-2 ^25^.

At CNB-CSIC, Vero E6 cell lines were kindly provided by Dr. Eric Snjider (University of Leiden Medical Center, The Netherlands). Both Huh7 and Vero E6 cell lines were cultured in Dulbecco-modified Eagle medium (DMEM) supplemented with 25 mM HEPES buffer, 2 mM l-glutamine (Sigma-Aldrich, St. Louis, MI, USA), 1% nonessential amino-acids (Sigma-Aldrich), 10% fetal bovine serum (FBS; BioWhittaker, Inc., Walkersville, MD, USA). In the post-infection semisolid medium, the percentage of FBS was reduced to 2%, and DEAE-dextran was added to a final concentration of 0.08 mg/mL.

At IRTA-CReSA, Vero E6 cells were obtained from the ATCC (ATCC CRL-1586) and cultured in DMEM (Lonza, Basel, Switzerland) supplemented with 5% (FBS (EuroClone, Pero, Italy), 100 U/mL penicillin, 100 μg/mL streptomycin, and 2 mM glutamine 8 (all ThermoFisher Scientific, Waltham, MA, USA). In the post-infection medium, the percentage of FBS was reduced to 2%.

### IgG ELISA testing procedures

Qualitative determination of IgG class antibodies cross-reactivity against antigens of the tested coronaviruses was performed using ELISA techniques. IVIG samples were serially diluted using the buffer solutions provided in each IgG ELISA kit. The following kits were used for the qualitative determination of IgG class antibodies in the experimental IVIG lots: SARS Coronavirus IgG ELISA kit (Creative Diagnostics, Shirley, NY, USA), against virus lysate; Human Anti-SARS-CoV-2 Virus Spike 1 [S1] IgG ELISA Kit (Alpha Diagnostic Intl. Inc., San Antonio, TX, USA), against S1 subunit spike protein; RV-402100-1, Human Anti-MERS-NP IgG ELISA Kit (Alpha Diagnostic Intl. Inc.), against N protein; RV-402400-1, Human Anti-MERS-RBD IgG ELISA Kit (Alpha Diagnostic Intl. Inc.), against receptor-binding domain (RBD) of S1 subunit spike protein (S1/RBD); RV-402300-1, Human Anti-MERS-S2 IgG ELISA Kit (Alpha Diagnostic Intl. Inc.), against S2 subunit spike protein; RV-405200 (formerly RV-404100-1). In all cases the determinations were carried out following the manufacturer’s instructions. Reactivity was rated as negative if no reaction was observed with neat IVIG or positive if the lowest IVIG dilution demonstrated reactivity.

### Neutralization assay for SARS-CoV, SARS-CoV-2 (MAD6 isolate) and MERS-CoV

IVIG samples were serially diluted (factor 10 dilutions: 1:10^2^, 1:10^3^, 1:10^4^, and 1:10^5^) in Dulbecco’s Phosphate Buffered Saline (PBS; Gibco, Thermo Fisher Scientific, Waltham, MA, USA). Samples of each IVIG dilution were incubated for 1 h (37°C; 5% CO_2_) with 300 plaque forming units (PFU) of SARS-CoV, SARS-CoV-2 or MERS-CoV. Aliquots of 50 μL of each IVIG dilution-virus complex were added in duplicate to confluent monolayers of Vero E6 cells (for SARS-CoV and SARS-CoV-2) or Huh7 (for MERS-CoV), seeded in 12-well plates and incubated for 1 h (37°C; 5% CO_2_). After this adsorption time, the IgG-virus complex inoculum was removed, and a semi-solid overlay was added (DMEM 2% FBS + 0.6% agarose). Cells were incubated for 72 h at 37°C. The semi-solid medium was removed, and the cells were fixed with 10% neutral buffered formaldehyde (Sigma-Aldrich) for 1 hour at room temperature and stained with 0.2% aqueous gentian violet for 10 min, followed by plaque counting. The sensitivity threshold of the technique was 20 PFU per mL.

The neutralization potency of the IVIG products was expressed in two ways: 1) percent reduction in PFU calculated from the PFU count after neutralization by IVIG relative to initial PFU count inoculated onto the cells; and 2) plaque reduction neutralization test (PRNT_50_) value, calculated as the −log_10_ of the reciprocal of the highest IVIG dilution to reduce the number of plaques by 50% compared to the number of plaques without IVIG.

### Neutralization assay for SARS-CoV-2 (EPI_ISL_418268 isolate)

A fixed concentration of a SARS-CoV-2 stock (10^1.8^ TCID 50/mL, a concentration that achieves 50% cytopathic effect) was mixed with decreasing concentrations of the IVIG samples (range 1:10 to 1:5120), each mixture was incubated for 1 h at 37° C and added to Vero E6 cells. To assess potential plasma-induced cytotoxicity, Vero E6 cells were also cultured with the same decreasing concentrations of plasma in the absence of SARS-CoV-2. Uninfected cells and untreated virus infected cells were used as negative and positive infection controls, respectively. Plasma from a COVID-19 positive patient with a high half-maximal inhibitory concentration (IC_50_) was included as an active positive control (expressed as the −log_10_ of the reciprocal of the dilution). All the cultures were incubated at 37°C and 5% CO_2_ for 3 days.

Cytopathic or cytotoxic effects of the virus or plasma samples were measured at 3 days post infection, using the Cell Titer-Glo luminescent cell viability assay (Promega, Madison, WI, USA). Luminescence was measured in a Fluoroskan Ascent FL luminometer (ThermoFisher Scientific). Neutralization curves are shown as nonlinear regressions. IC_50_ values were determined from the fitted curves as the plasma dilutions that produced 50% neutralization. Details of the technique are available elsewhere ^21^.

## Results

### Cross-reactivity studies (ELISA binding assays)

IVIG products showed consistent reactivity to antigens of SARS-CoV (culture lysate) at 10-100 mg/mL IgG, SARS-CoV-2 (S1 subunit protein) at 100 μg/mL IgG, and MERS-CoV (N protein, S1 subunit/RHD protein and S2 subunit protein) at 50 μg/mL IgG (Table 1).

**Table 1.** Results of IgG reactivity against SARS-CoV, SARS-CoV-2, and MERS-CoV. Two independent assays were performed on each IVIG product lot by ELISA. Concentration denotes the least potent IVIG dilution with positive result.

| IVIG product (lots) | % IgG | Country of origin of the plasma | Virus and antigen/target |  |  |  |  |
| --- | --- | --- | --- | --- | --- | --- | --- |
|  |  |  | SARS-CoV culture lysate | SARS-CoV-2 S1 subunit | MERS-CoV |  |  |
|  |  |  |  |  | N protein | S1 subunit/RBD | S2 subunit |
| F1 | 5% | Germany | 50 mg/mL | 100 µg/mL | 50 µg/mL | 50 µg/mL | 50 µg/mL |
| F2 | 5% | Czech Republic | 10 mg/mL | 100 µg/mL | 50 µg/mL | 50 µg/mL | 50 µg/mL |
| F3 | 10% | USA | 100 mg/mL | 100 µg/mL | 50 µg/mL | 50 µg/mL | 50 µg/mL |
| F4 | 10% | Spain | 100 mg/mL | 100 µg/mL | 50 µg/mL | 50 µg/mL | 50 µg/mL |
| G1 | 10% | USA | 100 mg/mL | 100 µg/mL | 50 µg/mL | 50 µg/mL | 50 µg/mL |
| G2 | 10% | USA | 100 mg/mL | 100 µg/mL | 50 µg/mL | 50 µg/mL | 50 µg/mL |

### Neutralization studies of SARS-CoV

All the assayed IVIG preparations had neutralizing activity against SARS-CoV ranging from 39% to 61% (Figure 1). All 10% IgG IVIG preparations (F3, F4, G1, and G2) showed PRNT_50_ neutralization titers between 2.0 and 3.3, corresponding to 50-61% PFU reduction (Figures 1B, 1C). The highest PFU reductions, 59.3% and 61.9% (PRNT_50_ neutralization titers of 3.2 and 3.3), were observed with lots F4 and G1, respectively, at 1 and 0.1 mg/mL IgG (dilution factors 2 and 3). The F1 and F2 lots, (5% IgG) showed a lower neutralization capacity with PFU reductions of 39.5% and 43.3%, respectively (Figure 1A).

**Figure 1.**
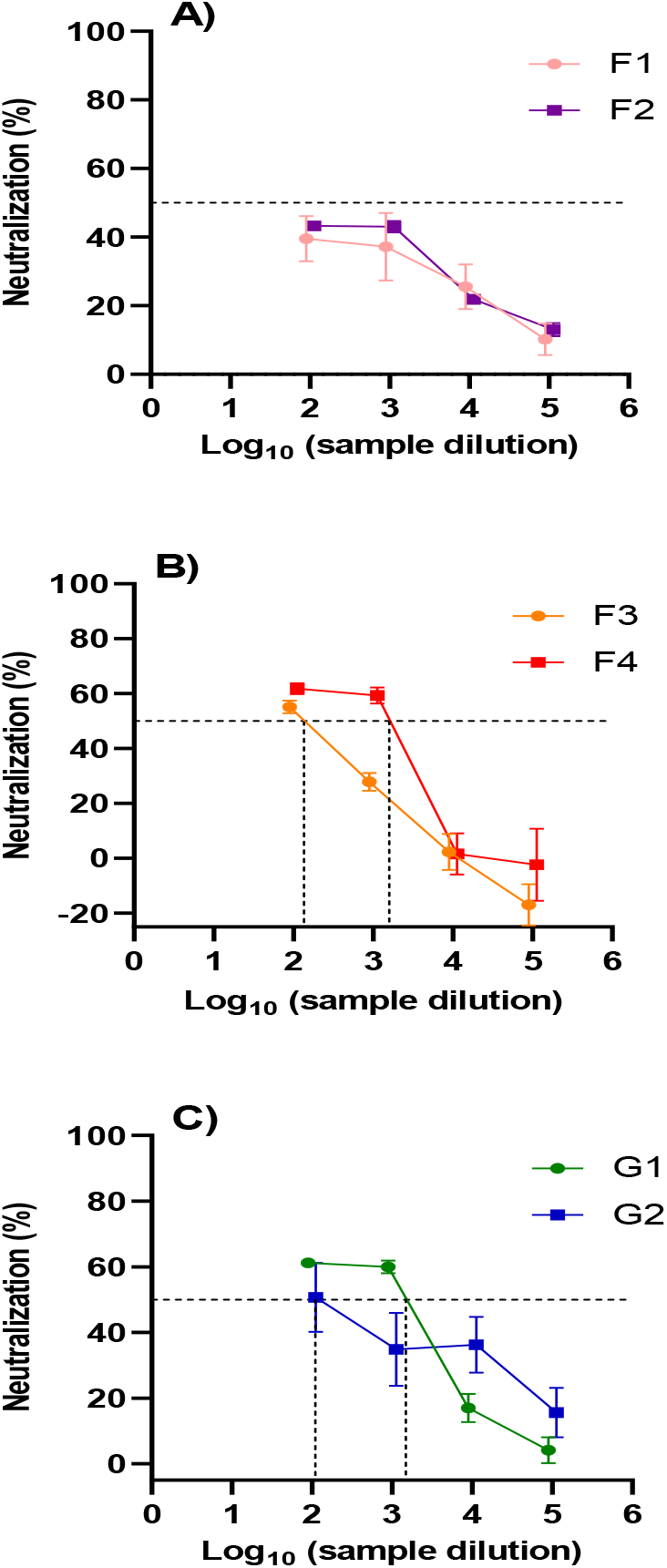
Cross-neutralization capacity of IVIG against SARS-CoV. The results were represented as percentage of neutralization calculated from reduction of PFU counts versus serial dilutions (1:102 – 1:105). The dotted lines indicate the PRNT_50_ values, i.e. the −log_10_ of the reciprocal of the highest IVIG dilution to reduce the number of plaques by 50%. Neutralization by: A) F1-F2 lots of Flebogamma 5% DIF; B) F3-F4 lots of Flebogamma 10% DIF; C) G1-G2 lots of Gamunex-C.

### Neutralization studies of SARS-CoV-2

For SARS-CoV-2 MAD6 isolate, all IVIG lots, except F1 (inconclusive results) showed a significant neutralizing activity and reached PRNT_50_ titers ranging from 4.5 to >5 (Figure 2). PFU reductions ranging from 78.2% to 82.5% were observed with lots F2, F3 and F4 at a dilution factor of 1. Even at the highest dilution factor (5 = 0.5 and 1 μg/mL), the PFU reduction ranged from 38.5% to 50.9% corresponding to PRNT_50_ titers of 4.5-5.0 (Figures 2A, 2B). For lots G1 and G2, the PFU reduction was even higher, ranging from 88.5% - 89.5% at a dilution factor of 1 to 61.7% −62.5% at a dilution factor of 5 with PRNT_50_ titers greater than 5 (Figure 2C).

**Figure 2.**
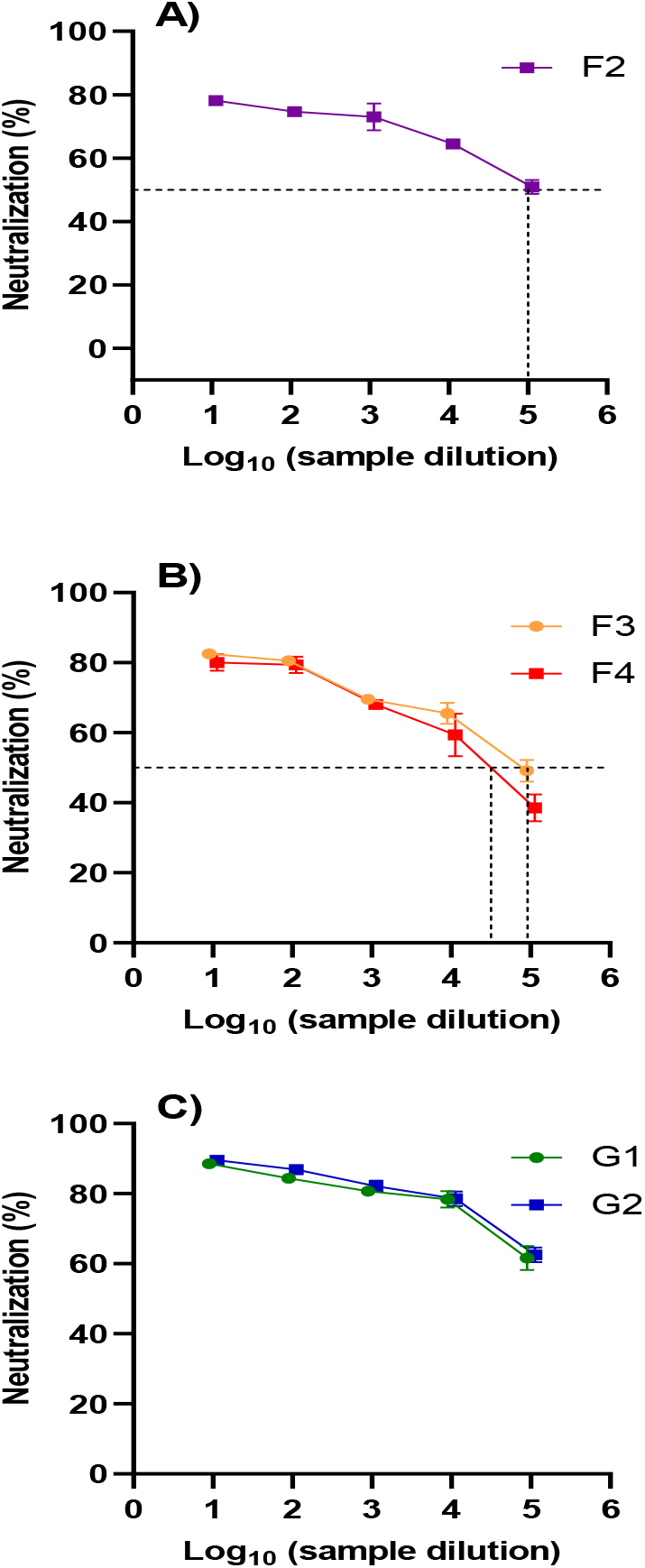
Cross-neutralization capacity of IVIG against SARS-CoV-2 (MAD6 isolate). The results were represented as percentage of neutralization calculated from reduction of PFU counts versus serial dilutions (1:102 – 1:105). The dotted lines indicate the PRNT_50_ values, i.e. the −log_10_ of the reciprocal of the highest IVIG dilution to reduce the number of plaques by 50%. Neutralization by: A) F1-F2 lots of Flebogamma 5% DIF; B) F3-F4 lots of Flebogamma 10% DIF; C) G1-G2 lots of Gamunex-C.

For the SARS-CoV-2 EPI_ISL_418268 isolate, F4 and G1 lots neutralized 58.4% and 64.7%, respectively, TCID_50_ counts at a dilution factor of 1 (Figure 3). As shown in Figure 3, one replicate of F4 product failed to demonstrate neutralization.

**Figure 3.**
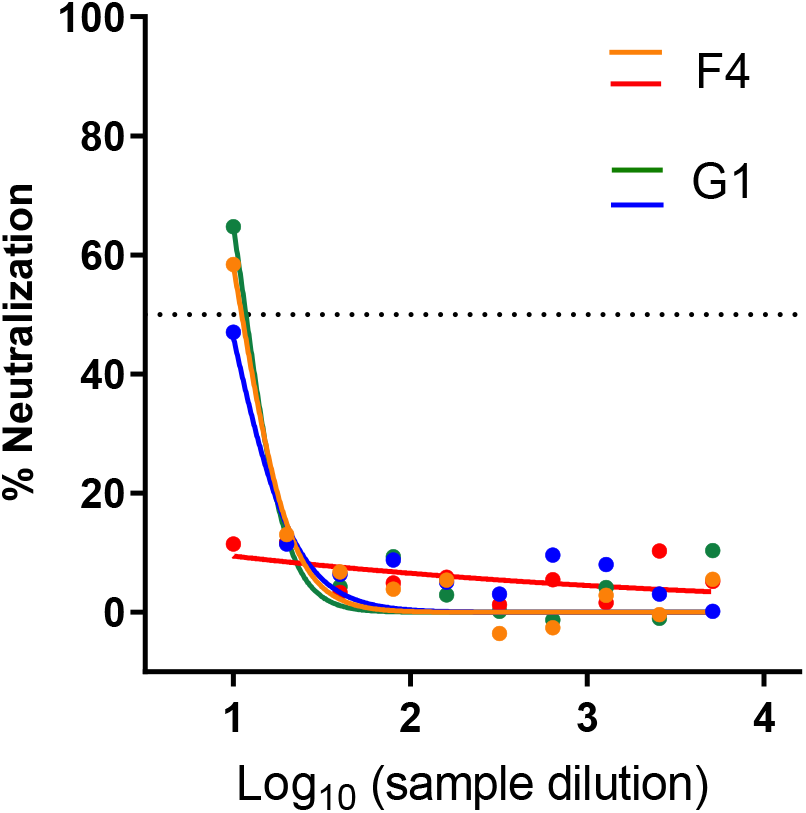
Cross-neutralization capacity of IVIG lots (F4: Flebogamma 10% DIF; G1: Gamunex-C) against SARS-CoV-2 (EPI_ISL_418268 isolate): The graphs represent the percent neutralization calculated by reduction of cytopathic effect versus serial dilutions.

### Neutralization studies of MERS-CoV

No IVIG lot showed any significant PFU reduction (i.e., >10%) on MERS-CoV even at the lowest dilution factor (10 mg/mL IgG).

## Discussion

The results presented here demonstrate, for the first time, significant cross-neutralization activity against SARS-CoV and especially SARS-CoV-2 in therapeutic IVIG concentrates (Flebogamma DIF and Gamunex-C). This neutralizing activity correlates with the cross-reactivity to different coronavirus antigens observed in ELISA binding assays with IVIG, as shown in a previous study ^19^. The plasma used to manufacture the tested IVIG lots was collected prior the detection of SARS-CoV-2 in Europe and the USA. Therefore, these results should be ascribed to cross-reactivity against SARS-CoV-2 by antibodies against endemic human coronaviruses in the human population at large. Similar results have been reported for SARS-CoV and MERS-CoV. ^16–18^

These neutralization studies showed that IVIG products contain antibodies with cross-neutralizing capacity against SARS-CoV (40-60%) and SARS-CoV-2 (80%-90%), but not against MERS-CoV (<10%). These results suggest that the cross-neutralizing antibodies target antigenic regions more conserved in SARS-CoV and SARS-CoV-2 than in MERS-CoV.

No significant differences in the neutralizing capacity were observed among IVIG lots regardless the country of origin for the plasma. This reinforces the broad applicability of these results. Two different neutralization techniques were used for SARS-CoV-2 and both techniques showed not only the IVIG neutralization capacity, but also the reliability of the results. In addition, results obtained with two different SARS-CoV-2 isolates confirm that the neutralization capacity is not dependent on the isolate. This was not unexpected since no significant sequence differences have been observed among SARS-CoV-2 isolates currently circulating throughout the world.

The percentage of SARS-CoV-2 cross-neutralization was higher in the PFU reduction technique than in the cytopathic effect/cytotoxic technique with very low or negative values in some few cases (inconclusive for lot F1 by the PFU study, and cytopathic effect in one replicate of lot F4). This suggests that the technique used and/or slight variations in methodology may significantly influence the nature or magnitude of the results. Therefore, further evaluation this cross-neutralizing activity should be conducted.

Cross-neutralization is gaining attention as a protective mechanism against viral infection in the context of the COVID-19 health emergency. The results of this study are in agreement with recent studies that describe cross-neutralization of SARS-CoV-2 by monoclonal antibodies from memory B cells of an individual who was infected with SARS-CoV ^26^. Furthermore, SARS-CoV-2-reactive CD4+ T cells have been detected in around half of unexposed individuals, suggesting that there is cross-reactive T cell recognition between circulating common cold coronaviruses and SARS-CoV-2 ^27^. However, the levels of cross-neutralizing antibodies against SARS-CoV-2 in the sera of SARS-CoV patients can be highly variable ^28^. IVIG products are prepared using plasma from thousands of different donors, hence containing a broad representation of the state of immunity in the population at that time. This is consistent with the low rate of variability found among the different lots of IVIG products tested.

Nevertheless, greater variability is expected among individuals with respect to infection by a given endemic human coronavirus. Therefore, it has been hypothesized that the diversity of symptoms observed in SARS-CoV-2-infected individuals and even the potential for getting infected may depend on pre-existing cross-immunity due to previous exposure to other endemic human coronaviruses. In this regard, a detailed study of the state of immunity in the general population distinguishing those affected and not affected by the SARS-CoV-2 may be warranted.

The higher cross-neutralizing capacity of the tested IVIG preparations against SARS-CoV and SARS-CoV-2 than MERS-CoV may be explained by higher sequence identity of the S proteins of circulating mild human coronaviruses (HCoV-OC43 and HCoV-HKU1) and SARS-CoV and SARS-CoV-2 compared to MERS-CoV (32%-33% vs. 23%-25) ^14,15^. Additionally, differences in specific domains of the S protein between SARS-CoV and SARS-CoV-2 might explain higher cross-reactivity of the tested IVIG against SARS-CoV-2 compared to SARS-CoV (80%-90% vs. 40%-60%). The absence of cross-neutralization against MERS-CoV despite the cross-reactivity observed in ELISA assays suggest that these antibodies are not neutralizing. This does not necessarily indicate that the antibodies are not functional by another mechanism. For example, these non-neutralizing antibodies could be labelling the virion for identification by immune cells and subsequent destruction ^29^.

Despite the limitations of the *in vitro* nature of this study, the clinical implications of the findings are encouraging. Although IVIG are considered a therapeutic option for hyperinflammation in patients with severe COVID-19 ^30^, the results of this study may support the use of high dose IVIG as a therapy for COVID-19. Positive results have already been reported for IVIG in case studies ^31,32^. IVIG is being tested in an ongoing clinical trial ^33^. Further studies looking at the functionality of these antibodies could improve our understanding the human coronavirus acquired immunity. This could pave the way for IVIG (and other IgG products such as intramuscular or subcutaneous preparations) as a potential therapeutic/prophylactic approach to fight future epidemics by emerging human coronaviruses.

In conclusion, under the experimental conditions of this study, IVIG (Flebogamma DIF and Gamunex-C) contained antibodies with significant neutralization capacity against SARS-CoV and SARS-CoV-2, but not against MERS-CoV. Additional research is warranted to advance IVIG towards clinical use for COVID-19.

## Acknowledgements

Jordi Bozzo PhD, CMPP and Michael K. James PhD (Grifols) are acknowledged for medical writing and editorial support in the preparation of this manuscript. Contribution from Antonio Páez MD (Grifols) who provided his expert opinion is acknowledged. The authors acknowledge the expert technical assistance from Daniel Casals, Eduard Sala, Judith Luque, Laura Gómez, and Gonzalo Mercado (Grifols, Viral and Cell Culture Laboratory).

## Disclosures

Neutralization experiments were funded by Grifols, the manufacturer of Flebogamma^®^ DIF and Gamunex^®^-C. JVA, JR, MBP, JMH, IS, and LE declare having no other conflict of interest. JMD, CR and RG are full-time employees of Grifols.

